# Microbiota profiling with long amplicons using Nanopore sequencing: full-length 16S rRNA gene and whole rrn operon

**DOI:** 10.1101/450734

**Authors:** Anna Cuscó, Carlotta Catozzi, Joaquim Viñes, Armand Sánchez, Olga Francino

## Abstract

**Background:** Profiling microbiome on low biomass samples is challenging for metagenomics since these samples are prone to present DNA from other sources, such as the host or the environment. The usual approach is sequencing specific hypervariable regions of the *16S rRNA* gene, which fails to assign taxonomy to genus and species level. Here, we aim to assess long-amplicon PCR-based approaches for assigning taxonomy at the genus and species level. We use Nanopore sequencing with two different markers: full-length *16S rRNA* (∼1,500 bp) and the whole *rrn* operon (*16S rRNA* gene – ITS – *23S rRNA* gene; 4,500 bp).

**Methods:** We sequenced a clinical isolate of *Staphylococcus pseudintermedius*, two mock communities (HM-783D, Bei Resources; D6306, ZymoBIOMICS™) and two pools of low-biomass samples (dog skin). Nanopore sequencing was performed on MinION™ (Oxford Nanopore Technologies) using 1D PCR barcoding kit. Sequences were pre-processed, and data were analyzed using WIMP workflow on EPI2ME (ONT) or Minimap2 software with *rrn* database.

**Results:** Full-length *16S rRNA* and the *rrn* operon retrieved the microbiota composition from the bacterial isolate, the mock communities and the complex skin samples, even at the genus and species level. For *Staphylococcus pseudintermedius* isolate, when using EPI2ME, the amplicons were assigned to the correct bacterial species in ∼98% of the cases with *rrn* operon as the marker, and ∼68% of the cases with *16S rRNA* gene respectively. In both skin microbiota samples, we detected many species with an environmental origin. In chin, we found different *Pseudomonas* species in high abundance, whereas in dorsal skin there were more taxa with lower abundances.

**Conclusions:** Both full-length *16S rRNA* and the *rrn* operon retrieved the microbiota composition of simple and complex microbial communities, even from the low-biomass samples such as dog skin. For an increased resolution at the species level, *rrn* operon would be the best choice.

## Introduction

The microbiota profile of low biomass samples such as skin is challenging for metagenomics. These samples are prone to present DNA contamination from the host or exogenous sources, which can overcome the DNA of interest [1, 2]. Thus, the usual approach is amplifying and sequencing certain genetic markers that are ubiquitously found within the studied kingdom rather than performing metagenomics. Ribosomal marker genes are a common choice: *16S rRNA* and *23S rRNA* genes for taxonomically classify bacteria [3, 4]; ITS1, and ITS2 regions for fungi [5, 6].

Until now, most of the microbiota studies rely on second-generation sequencing (massive parallel sequencing), and target a short fragment of the *16S rRNA* gene, which presents nine hypervariable regions (V1-V9) that are used to infer taxonomy [7, 8]. The most common choices for host-associated microbiota are V4 or V1-V2 regions that present different taxonomic coverage and resolution depending on the taxa [9]. V4 region represents better the whole bacterial diversity, although it fails to amplify *Cutibacterium acnes* (formerly called *Propionibacterium acnes)* that is a ubiquitous skin commensal in humans. So, when performing a skin microbiota study, the preferred choice is V1-V2 regions, although they lack sensitivity for the *Bifidobacterium* genus and it poorly amplifies the Verrucomicrobia phylum [10].

Apart from the biases derived from the primer choice, short fragment strategies usually fail to assign taxonomy reliably down to genus and species level. This taxonomic resolution is particularly useful when associating microbiota to clinics such as in characterizing disease status or when developing microbiota-based products, such as pre- or pro-biotics [11]. For example, in human atopic dermatitis (AD) the signature for AD-prone skin when compared to healthy skin was enriched for *Streptococcus* and *Gemella* but depleted for *Dermacoccus*. Moreover, nine different bacterial species were identified to have significant AD-associated microbiome differences [12]. In canine atopic dermatitis, *Staphylococcus pseudintermedius* has been classically associated with the disease. Microbiota studies of canine atopic dermatitis presented an overrepresentation of *Staphylococcus* genus [13, 14], but the species was confirmed when complementing the studies using directed qPCRs for the species of interest [13] or using a *Staphylococcus*-specific database and V1-V3 region amplification [14].

With the launching of third-generation single-molecule technology sequencers, these short-length associated issues can be overcome by sequencing the full-length of *16S rRNA* gene (1,500 bp) or even the whole *rrn* operon (4,500 bp) that includes *16S rRNA* gene, ITS region, and 23S rRNA gene. MinION^TM^ sequencer of Oxford Nanopore Technologies (ONT) is a single-molecule sequencer that is portable, affordable with a small budget and offers long-read output. Its main limitation is the still high error rate.

Several studies targeting the full-length *16S rRNA* gene have already been performed using nanopore sequencing to: i) characterize artificial and already characterized bacterial communities (mock community) [15–17]; ii) characterize complex microbiota samples, from mouse gut [18], wastewater [19], microalgae [20] and dog skin [21]; and iii) characterize the pathogenic agent in a clinical sample [22–24]. On the other hand, only two studies have been performed using the whole *rrn* operon to characterize mock communities [25] and complex natural communities [26].

Here we aim to assess the potential of Nanopore sequencing using both the full-length *16S rRNA* (1,500bp) and the whole *rrn* operon (4,500bp) in: i) a clinical isolate of *Staphylococcus pseudintermedius*, ii) two bacterial mock communities; and iii) two complex skin microbiota samples.

## Material and methods

### Samples and DNA extraction

We used two DNA mock communities as simple, well-defined microbiota samples:

- HM-783D, kindly donated by BEI resources (http://www.beiresources.org) that contained genomic DNA from 20 bacterial strains with staggered ribosomal RNA operon counts (from 1,000 to 1,000,000 copies per organism per μL).
- ZymoBIOMICS™ Microbial Community DNA (http://www.zymoresearch.com) Standard that contained a mixture of genomic DNA extracted from pure cultures of eight bacterial strains.

We have also sequenced a pure bacterial isolate of *Staphylococcus pseudintermedius* obtained from an ear of a dog affected with otitis.

As a complex microbial community, we used two DNA sample pools from skin microbiota of healthy dogs targeting two different skin sites: i) dorsal back: DNA from two dorsal samples from Beagle dogs; and ii) chin: DNA from five chin samples from Golden-Labrador Retriever crossed dogs. Skin microbiota samples were collected using Sterile Catch-All™ Sample Collection Swabs (Epicentre Biotechnologies) soaked in sterile SCF-1 solution (50 mM Tris buffer (pH = 8), 1 mM EDTA, and 0.5% Tween-20). DNA was extracted from the swabs using the PowerSoil™ DNA isolation kit (MO BIO) and blank samples were processed simultaneously (for further details on sample collection and DNA extraction see [27]).

### PCR amplification of ribosomal markers

Two ribosomal markers were evaluated in this study: full-length *16S rRNA* gene (∼1,500 bp) and the whole *rrn* operon (∼4,500 bp). Before sequencing, bacterial DNA was amplified using a nested PCR, with a first PCR to add the specific primer sets (Table 1) tagged with the Oxford Nanopore universal tag and a second PCR to add the barcodes from the barcoding kit (EXPPBC001). Each PCR reaction included a no template control (NTC) sample to assess possible reagent contamination.

**Table 1.**
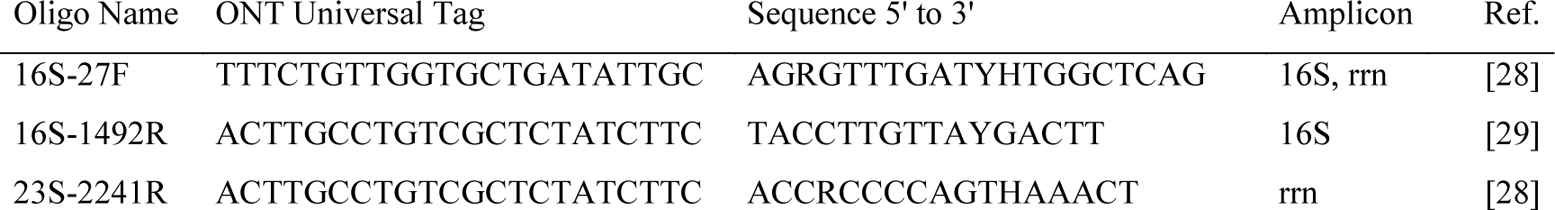
Primer sequences for full-length *16S rRNA* gene and *rrn* operon amplification

For the first PCR, we targeted the full *16S rRNA* gene using 16S-27F and 16S-1492R primer set and the whole *rrn* operon (*16S rRNA* gene – ITS – *23S rRNA* gene) using 16S-27F and 23S-2241R primer set (Table 1).

PCR mixture for full-length *16S rRNA* gene (25 μl total volume) contained 5 ng of DNA template, 5 μl of 5X Phusion^®^ High Fidelity Buffer, 2.5 μl of dNTPs (2 mM), 1 μl of 16S-27F (0.4 μM), 2 μl of 16S-1492R (0.8 μM) and 0.25 μl of Phusion^®^ Hot Start II Taq Polymerase (0.5 U) (Thermo Scientific, Vilnius, Lithuania). The PCR thermal profile consisted of an initial denaturation of 30 s at 98 °C, followed by 25 cycles of 15 s at 98 °C, 15 s at 51 °C, 45 s at 72 °C, and a final step of 7 min at 72 °C.

PCR mixture for *rrn* whole operon (50 μl total volume) contained 5 ng of DNA template, 10 μl of 5X Phusion^®^ High Fidelity Buffer, 5 μl of dNTPs (2 mM), 5 μl of each primer (1 μM) and 0.5 μl of Phusion^®^ Hot Start II Taq Polymerase (1 U). The PCR thermal profile consisted of an initial denaturation of 30 s at 98 °C, followed by 25 cycles of 7 s at 98 °C, 30 s at 59 °C, 150 s at 72 °C, and a final step of 10 min at 72 °C.

The amplicons were cleaned-up with the AMPure XP beads (Beckman Coulter) using a 0.5X and 0.45X ratio for the *16S rRNA* gene and the whole *rrn* operon, respectively. Then they were quantified using Qubit™ fluorometer (Life Technologies, Carlsbad, CA) and volume was adjusted to begin the second PCR with 0.5 nM of the first PCR product or the whole volume when not reaching the required concentration (mostly for samples that amplified the *rrn* operon).

PCR mixture for the barcoding PCR (100 μl total volume) contained 0.5 nM of first PCR product, 20 μl of 5X Phusion^®^ High Fidelity Buffer, 10 μl of dNTPs (2 mM), and 1 μL of Phusion^®^ Hot Start II Taq Polymerase (2 U). Each PCR tube contained the DNA, the PCR mixture and 2 μl of the specific barcode. The PCR thermal profile consisted of an initial denaturation of 30 s at 98 °C, followed by 15 cycles of 7 s at 98 °C, 15 s at 62 °C, 45 s (for *16S rRNA* gene) or 150 s (for *rrn* operon) at 72 °C, and a final step of 10 min at 72 °C.

Again, the amplicons were cleaned-up with the AMPure XP beads (Beckman Coulter) using a 0.5X and 0.45X ratio for the *16S rRNA* gene and the whole *rrn* operon, respectively. For each sample, quality and quantity were assessed using Nanodrop and Qubit™ fluorometer (Life Technologies, Carlsbad, CA), respectively.

In most cases, the different barcoded samples were pooled in equimolar ratio to obtain a final pool (1000-1500 ng in 45 μl) to do the sequencing library.

### Nanopore sequencing library preparation

The Ligation Sequencing Kit 1D (SQK-LSK108 by ONT) was used to prepare the amplicon library to load into the MinION^TM^, following the instructions of the 1D PCR barcoding amplicon protocol of ONT. Input DNA samples were composed of 1-1.5 μg of the barcoded DNA pool in a volume of 45 μL and 5 μL of DNA CS (DNA from lambda phage, used as a positive control in the sequencing). The DNA was processed for end repair and dA-tailing using the NEBNext End Repair / dA-tailing Module (New England Biolabs). A purification step using 1X Agencourt AMPure XP beads (Beckman Coulter) was performed.

For the adapter ligation step, a total of 0.2 pmol of the end-prepped DNA were added in a mix containing 50 μL of Blunt/TA ligase master mix (New England Biolabs) and 20 μL of adapter mix and then incubated at room temperature for 10 min. We performed a purification step using Adapter Bead Binding buffer (provided on SQK-LSK108 kit) and 0.5X Agencourt AMPure XP beads (Beckman Coulter) to finally obtain the DNA library.

We prepared the pre-sequencing mix (14 μL of DNA library) to be loaded by mixing it with Library Loading beads (25.5 μL) and Running Buffer with fuel mix (35.5 μL). We used two SpotON Flow Cells Mk I (R9.4.1) (FLO-MIN106). After the quality control, we primed the flowcell with a mixture of Running Buffer with fuel mix (RBF from SQK-LSK108) and Nuclease-free water (575 μL + 625 μL). Immediately after priming, the nanopore sequencing library was loaded in a dropwise fashion using the SpotON port.

Once the library was loaded, we initiated a standard 48h sequencing protocol using the MinKNOW™ software.

### Data analysis workflow

The samples were run using the MinKNOWN software. After the run, fast5 files were basecalled and demultiplexed using Albacore v2.3.1. A second demultiplexing round was performed with Porechop [30], where only the barcodes that agreed with Albacore were kept. Porechop was also used to trim the barcodes and the adapters from the sequences (Figure 1). Moreover, we removed 45 extra basepairs from each end that correspond to the length of the universal tags and custom primers. After the trimming, reads were selected by size: 1,200 bp to 1,800 bp for *16S rRNA* gene; and 3,500 to 5,000 bp for rrn operon. We mapped the sequences obtained to the *rrn* database using Minimap2 [31]. Afterwards chimeras were detected and removed using yacrd [32].

**Figure 1.**
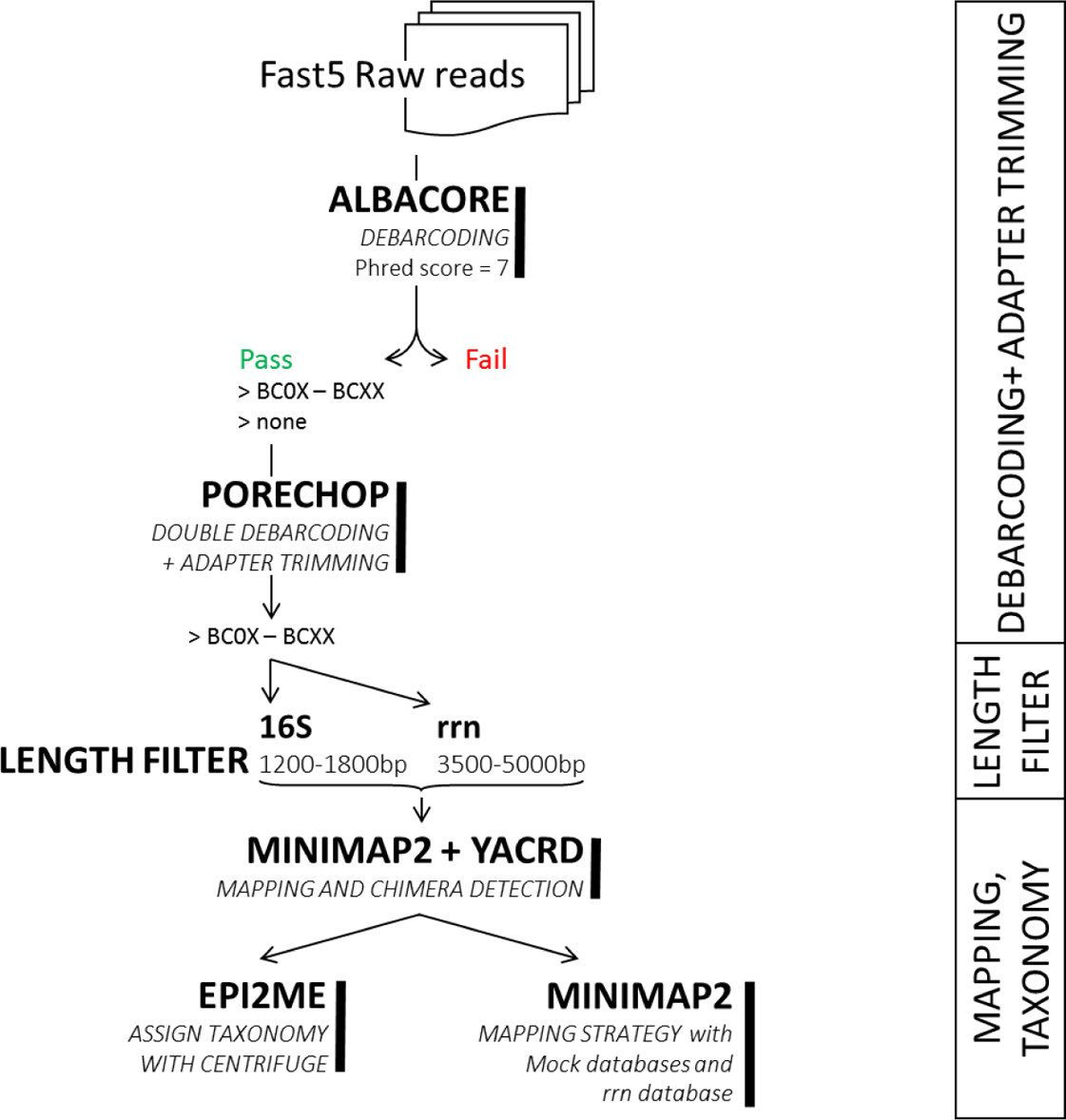
Bioinformatics analysis workflow from raw reads to final data.

To assign taxonomy to the trimmed and filtered reads we used to strategies: 1) a mapping-based strategy using Minimap2 [31]; or 2) a taxonomic classifier using What’s in my pot (WIMP) [33], a workflow from EPI2ME in ONT cloud (based on Centrifuge software [34]).

For the mapping-based strategy, we performed Minimap2 again with the non-chimeric sequences. We applied extra filtering steps to retain the final results: we kept only those reads that aligned to the reference with a block larger than 1,000 bp (for *16S rRNA* gene) and 3,000 bp (for whole *rrn* operon). For reads that hit two or more references, only the alignments with the highest alignment score (Smith-Waterman alignment score) were kept. After these filtering, the multimapping was mostly present in cases with entries that belonged to the same taxonomy.

The reference databases used in this study were:

- Mock database: a collection of the complete genomes that were included in each mock community, as described by the manufacturer. The HM-783D database was retrieved from NCBI using the reference accession numbers, while Zymobiomics mock community has already its database online on the Amazon AWS server.
- *rrn* database: sequences from the whole operon retrieved from Genbank [25].

For the taxonomic classification using *What’s In My Pot* (WIMP) workflow from Oxford Nanopore that uses NCBI database, only those hits with a classification score >300 were kept [34].

## Results

### Quality filtering results

After Albacore basecalling and Porechop processing, we lost around 5% of the initial reads (13% - 3%). After length trimming step, we lost more sequences (Table 1). In general, the samples amplified using *16S rRNA* marker gene recovered a higher percentage of reads after the quality control when compared to rrn operon: 74 - 95% *vs.* 32 - 80%. Especially for *rrn* operon, the largest percentage of reads was lost during the length trimming step: some of the reads included in that barcode presented the length of the *16S rRNA* gene.

After this first quality control, we performed an alignment with the mock and the *rrn* databases and checked for chimeras. Chimeras detected were dependent on the database used for the alignment. As a positive control, we used mock samples with their mock database. Chimera ratio was higher for *16S rRNA* gene amplicons (around ∼40%) than for *rrn* operon (∼10%), suggesting that PCR conditions for *16S rRNA* gene need to be adjusted or the PCR cycles reduced.

To conclude, the final useful sequences when amplifying for either amplicon were ∼40%. In *16S rRNA* gene, sequences were lost in chimera checking step. In *rrn* operon, sequences were lost in the length trimming step, probably due to the underrepresentation of the amplicon in the flowcell, since we ran them together with full-length *16S rRNA* amplicons in the same flow-cell.

### Mock community analyses

Microbial Mock Community HM-783D contained genomic DNA from 20 bacterial strains with staggered ribosomal RNA operon counts (from 1,000 to 1,000,000 copies per organism per µL). The bacterial composition detected should be proportional to the operon counts. This mock community would allow us determining if our approach reliably represents the actual bacterial composition of the community, especially considering low-abundant species.

We analyzed HM-783D mock community against its own database, which contains only the 20 representative species. On the one hand, using *16S rRNA* gene we were able to detect all the bacterial species present in the mock community, even the low-abundant ones. On the other hand, using the *rrn* operon we were able to detect only the most abundant species (at least 10^4^ operon copies) (Figure 2). This could be due to the lower sequencing depth obtained with *rrn* when compared with *16S rRNA,* and probably due to the underrepresentation of the *rrn* amplicon in the flowcell when running together with the full-length *16S rRNA* amplicons in the same flow-cell, as detailed above. Moreover, the relative abundances of *rrn* operon sequences were more biased than those obtained from *16S rRNA* gene sequencing, when compared to those expected, which confirmed that the primers for *rrn* need to be improved for universality.

**Figure 2.**
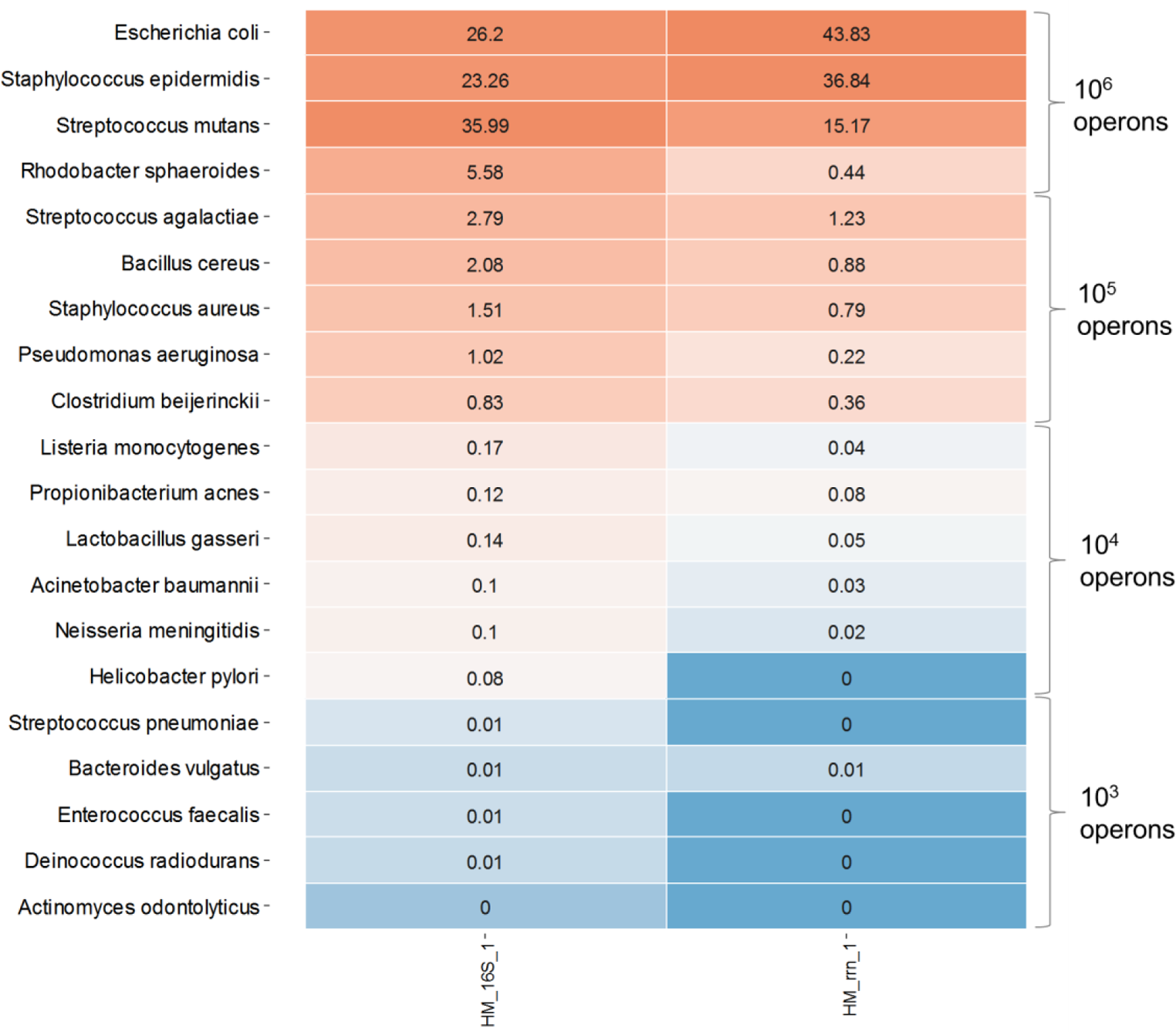
Heatmap representing the HM-783D mock community when mapped with its mock database. The darkest blue represents the bacteria that were not detected (< 10^4^ copies with *rrn* operon).

Zymobiomics mock community presents the same amount of genomic DNA from 8 different bacterial species; the expected *16S rRNA* gene content for each representative is also known, so we are able to determine if our approach represents the actual bacterial composition of the community reliably.

Both *16S rRNA* gene and *rrn* operon sequencing were able to detect 8 out of 8 bacterial species for Zymobiomics mock community, using Minimap2 and WIMP. The “Other taxa” group in Figure 3A can indicate: 1) not expected taxa (wrongly-assigned species, or previous contamination); or 2) higher taxonomic rank taxa (sequences not assigned to species level).

**Figure 3.**
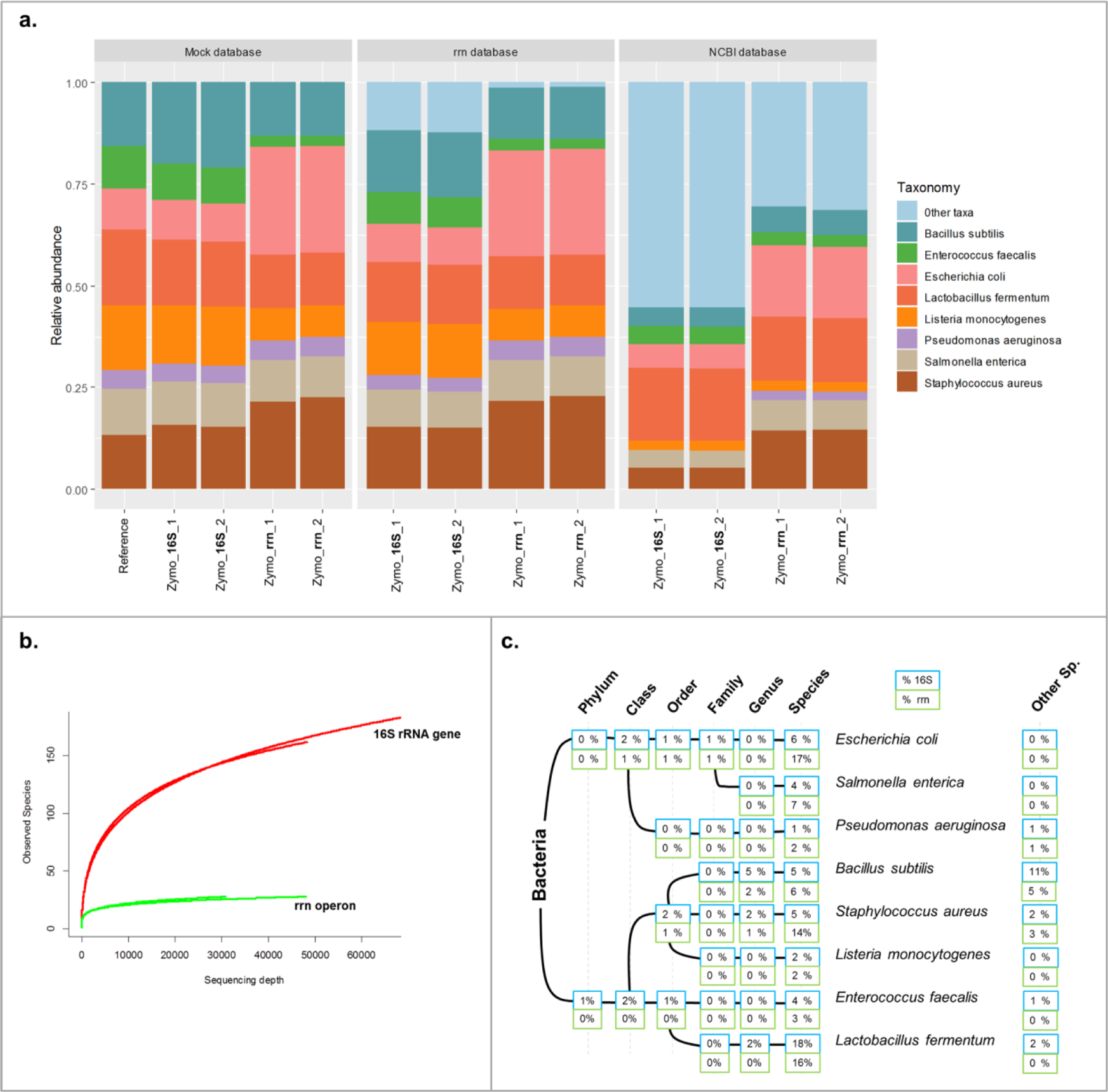
Zymobiomics mock community taxonomic analysis and diversity. A) Bar plots representing the relative abundance of the Zymobiomics mock community. “REF” bar represents the theoretical composition of the mock community regarding the *16S rRNA* gene content of each bacterium. B) Alpha diversity rarefaction plot using the rrn database at the species level. C) WIMP output represented in a taxonomic tree and percentage of reads in each taxonomic rank.

Using the mock community database (that contains only the 8 members of that community), we aimed to assess the biases regarding the actual abundance profile. *16S rRNA* gene better represented the bacterial composition of the mock community, when considering the abundances. The *rrn* operon amplification over-represented *Escherichia coli* and *Staphylococcus aureus* and under-represented *Enterococcus faecalis*.

The *rrn* database [25] contains 22,351 different bacterial species, including representatives of the species in the mock community. When using the *rrn* database, we found that the *rrn* operon was a better marker than *16S rRNA*: more than 98% of the sequences mapped to the corresponding species, and only < 2% of the total sequences mapped to a wrong species with *rrn* operon, whereas ∼15% of the sequences hit a wrong taxonomy with *16S rRNA*. We performed alpha diversity analyses using the same *rrn* database. The *rrn* operon hit 26 different species, whereas *16S rRNA* over-estimated the actual diversity, with hits to 202 different species (Figure 3B). However, when considering abundances, the diversity values are more similar presenting a Shannon index of 1.95 and 2.51, when using rrn operon and *16S rRNA* respectively (at 30,000 sequences/sample).

Using WIMP, we confirmed again the higher resolution power of *rrn* operon: ∼70% of the sequences were assigned to the correct species compared to ∼45% for *16S rRNA* gene. Among all the bacterial species included in the mock community, *Bacillus subtilis* presented more trouble for the correct taxonomic classification. The theoretically expected abundance for *B. subtilis* is 17% using *16S rRNA* gene. When using WIMP, only 5% of the total sequences were correctly classified at the species level, another 5% was classified correctly at the genus level, and another 10% was incorrectly classified as other *Bacillus* species (Figure 3C).

Apart from the mock communities, we also sequenced an isolate of *Staphylococcus pseudintermedius* obtained from canine otitis. When using WIMP approach with *rrn* operon, 97.5 % of the sequences were correctly assigned to the *S. pseudintermedius*. However, with *16S rRNA* gene, 68 % of the sequences were correctly assigned at the species level and 13% at the genus (Table 3). The wrong assigned species for *rrn* operon was ∼ 2.5 %, compared to ∼ 20 % for *16S rRNA* gene. On the other hand, through mapping the sequences to *rrn* database using Minimap2, we obtained no hit to *S. pseudintermedius*, since there is no representative in the *rrn* database. Instead, they were hitting mostly to *Staphylococcus schleiferi,* which is a close species, fewer also to *Staphylococcus hyicus* and *Staphylococcus agnetis*. These results highlight the need of comprehensive databases that include representatives of all the microorganisms relevant to a microbiome to correctly assign taxonomy.

### Complex microbial community analyses

After the first analyses with the mock communities, we were able to detect that the taxonomic resolution was higher when using *rrn* operon; the abundance profile was more reliable using *16S rRNA* marker gene though. If a bacterial species is not present in the database, the mapping strategy will give us the closest sequence resulting to an inaccurate taxonomic profile, such as we have seen for the *Staphylococcus pseudintermedius* isolate.

Here, we aimed to taxonomically profile two complex and uncharacterized microbial communities from dog skin (chin and dorsal) using the two different markers and comparing the mapping strategy (Minimap2 and *rrn* database) with WIMP workflow (NCBI database).

For chin samples of healthy dogs, we found a high abundance of *Pseudomonas* species followed by other genus with lower abundances such as *Erwinia* and *Pantoea*. Focusing on *Pseudomonas*, and going down to species level we were able to detect that the most abundant species was *Pseudomonas koreensis*, followed by *Pseudomonas putida* and *Pseudomonas fluorescens* (Figure 4A and Supplementary Table S1). On the other hand, dorsal skin samples were dominated by bacteria from the genera *Stenotrophomonas*, *Sanguibacter,* and *Bacillus*. We reached species level for *Stenotrophomonas rhizophila* and *Sanguibacter keddieii*. It is worth to note that *Glutamicibacter arilaitensis* is the same species as *Arthrobacter arilaitensis* but with a newer nomenclature [35] (Figure 4B and Supplementary Table S1). For both skin samples replicates, the results of the most abundant species converged and allowed characterizing this complex low-biomass microbial community at the species level.

**Figure 4.**
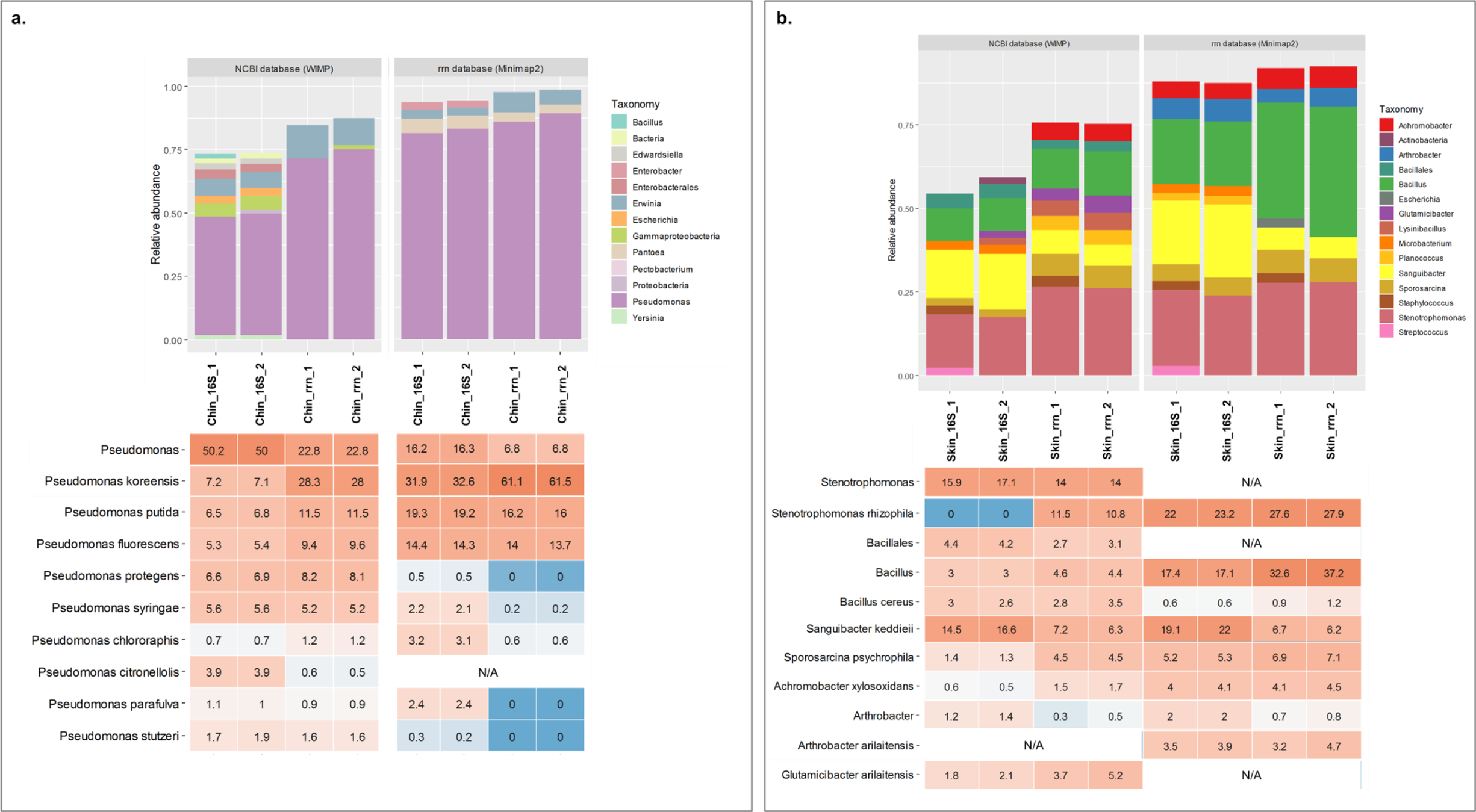
Microbiota composition of complex communities: skin samples of healthy dogs. A) Chin samples: upper part of the graphic, bar plot of the composition at the genus level using WIMP (left) and Minimap2 (right); lower part, heatmap of the *Pseudomonas* species within the community (scaled at 100%). B) Dorsal skin samples: upper part of the graphic, bar plot of the composition at the genus level using WIMP (left) and Minimap2 (right); lower part, heatmap of the ten most abundant taxa within the community. N/A in the heatmap indicates that taxon was not present in the database.

Finally, analyzing the dorsal skin samples, we also detected the presence of contamination from the previous nanopore run (Table 2). We sequenced dorsal skin samples twice: one with a barcode previously used for sequencing the HM-783D mock community and another one with a new barcode. We were able to detect mock community representatives within the re-used barcode (Figure 5). Some of them were found only in the sample that was using the re-used barcode (Sample_1); others were also present in the skin sample such as *Bacillus cereus* or *Staphylococcus aureus*. In total, this contamination from the previous run was representing ∼ 6% of the sample composition.

**Table 2.**
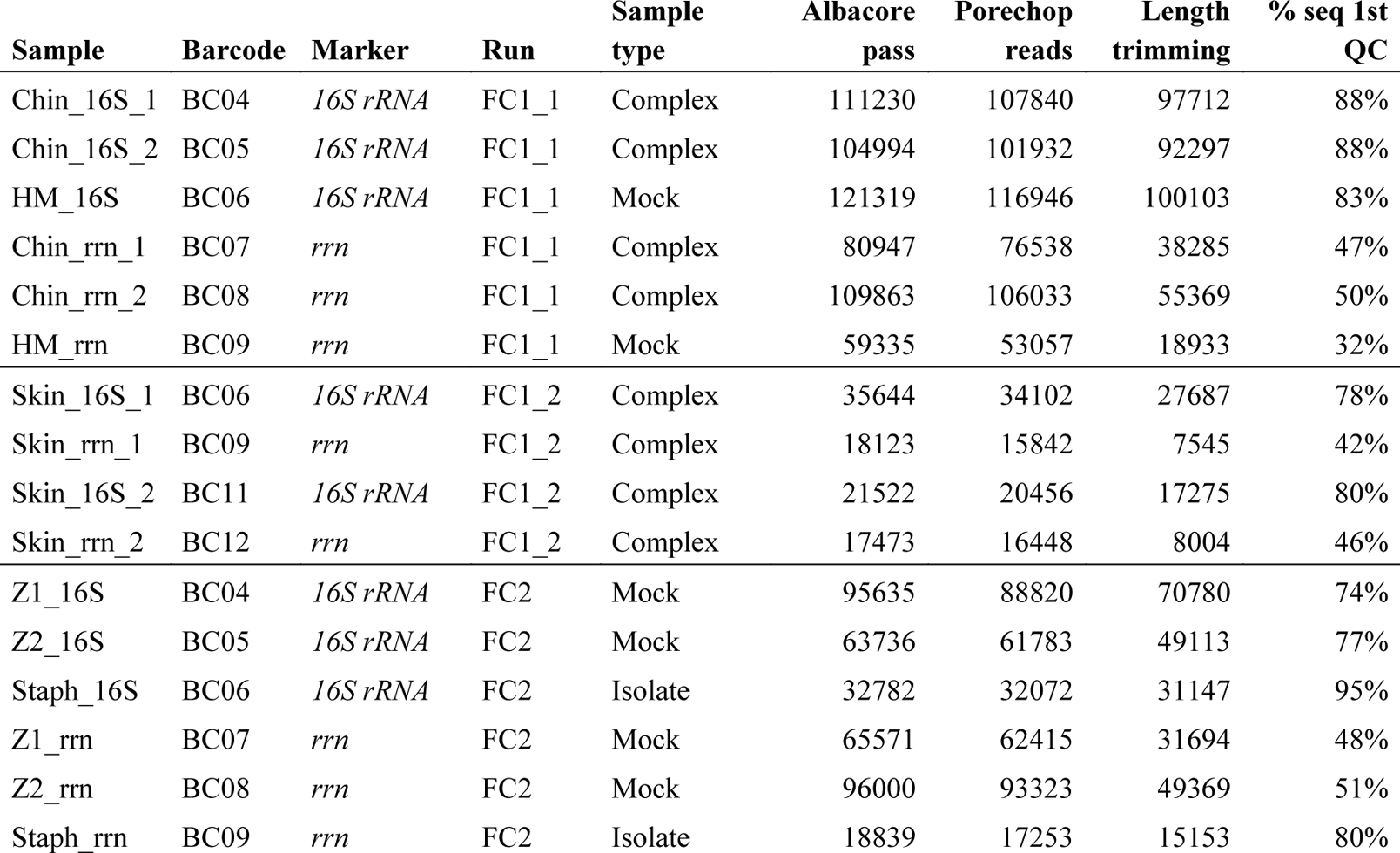
Samples included in the study and quality control results. HM - HM-783D mock community (BEI resources); Z1 and Z2 are replicates of ZymoBIOMICS™ Microbial Community DNA; Chin – microbiota from a pool of canine chin samples; Skin – microbiota from a pool of dorsal skin samples.

**Table 3.**
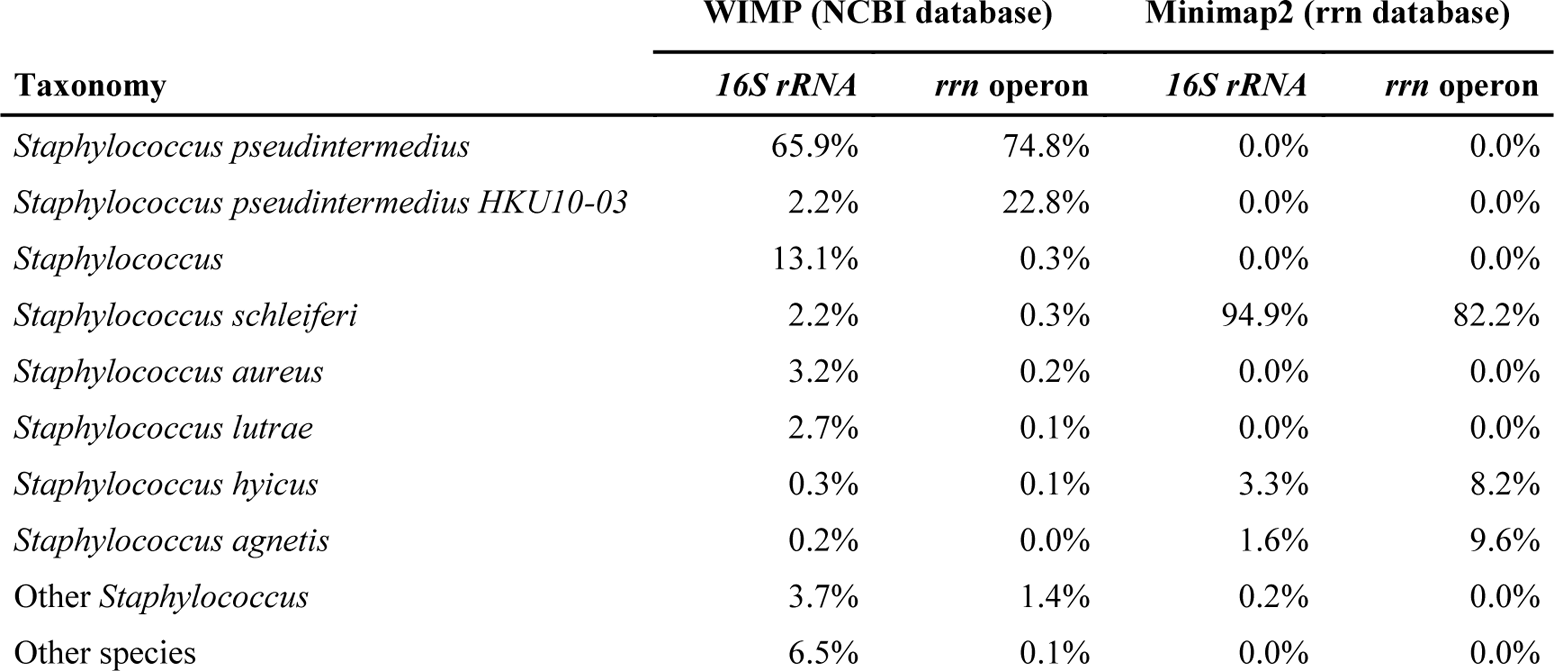
Taxonomy assignments of *S. pseudintermedius* isolate using WIMP workflow with NCBI database and Minimap2 with *rrn* database.

**Figure 5.**
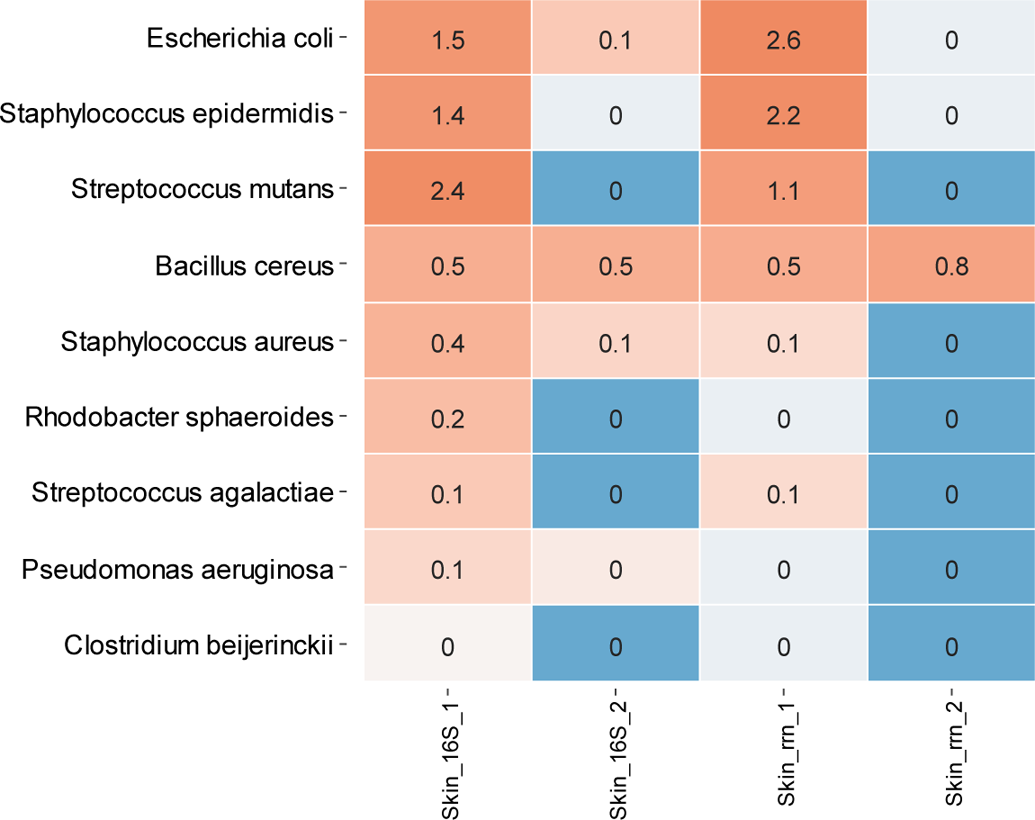
Heatmap representing the HM-783D mock community contamination. Samples Skin_1 are the ones reusing the HM-783D barcode from the previous run within the same flowcell. Samples Skin_2 are using a new barcode (not used in a prior run of the same flowcell). The darkest blue represents the bacteria that were not detected.

## Discussion

Full-length *16S rRNA* and the *rrn* operon retrieved the microbiota composition from the bacterial isolate, the mock communities and the complex skin samples, even at the genus and species level. Although Nanopore sequencing still presents a high error rate (average accuracy for the *S. pseudintermedius* isolate: 89%), we compensated this low accuracy with longer fragments to assess the taxonomy of several bacterial communities. In general, the longer the marker, the higher the taxonomical resolution both when using mapping software, such as Minimap2, or taxonomy classifiers such as WIMP in EPI2ME cloud.

When using EPI2ME (WIMP with NCBI database), the amplicons from the isolate *S. pseudintermedius* were assigned to the correct bacterial species in ∼98% and ∼68% of the cases, using *rrn* operon and *16S rRNA* operon respectively. In a previous study, Moon and collaborators used the full-length *16S rRNA* gene for characterizing an isolate of *Campylobacter fetus* and the marker assigned the species correctly for ∼89% of the sequences using EPI2ME [23]. The ratio of success on the correct assignment at species level depends on the species itself and its degree of sequence similarity in the selected marker gene. Within *Staphylococcus* genus, *16S rRNA* gene presents the highest similarity (around ∼97%) when compared to other genetic markers [36]. On the other hand, we observed that using the mapping strategy (through Minimap2) could lead to a wrong assigned species if the interrogated bacterium has not any representative on the chosen database. This strategy provides faster results than EPI2ME, but it needs an accurate comprehensive and representative database.

Analyses of the mock communities allowed us detecting if our approach represented the actual bacterial composition reliably, also when taking into account the low-abundant species (with the HM-783D staggered mock community). When using the *16S rRNA* marker gene, we were able to detect all bacterial members of both mock communities. However, when using *rrn* operon some of the low-abundant species were not detected. The likely reason is that we obtained lower number of reads for this marker, up to one magnitude. Mock communities also allowed us detecting the potential biases of our primer sets for both markers, since some of the species detected were over- and under-represented. *Actinomyces odontolyticus* and *Rhodobacter sphaeroides* seem to not amplify properly, neither with *16S rRNA* gene or rrn operon. Previous studies also detected the same pattern for these specific bacteria even when using or comparing different primer sets [16, 21]. Overall, *16S rRNA* primer set seemed less biased than *rrn* operon. When using the *rrn* operon, *E. coli* and *S. aureus* were overrepresented whereas others were underrepresented, suggesting that the primers should be improved for universality.

Focusing on dog chin samples, we could detect that mostly *Pseudomonas* species colonized them:*P. koreensis*, *P. putida,* and *P. fluorecens* as the main representatives. Recently, Meason-Smith and collaborators found *Pseudomonas* species associated with malodor in bloodhound dogs [37]. However, they were not the main bacterium found within the skin site tested but a low-abundant one, differing from what we have found here. On the other hand, Riggio and collaborators detected *Pseudomonas* as one of the main genera in canine oral microbiota in the normal, gingivitis and periodontitis groups [38]. However, the *Pseudomonas* species were not the same ones that we have detected here. It is worhty to notice that we had characterized these chin samples (and others) with 16S V1-V2 amplicons in a previous study [27], where we found some mutual exclusion patterns for *Pseudomonadaceae* family. This taxon showed an apparently “invasive pattern”, which could be mainly explained for the recent contact of the dog with an environmental source that contained larger bacterial loads before sampling [27]. Thus, our main hypothesis is that the *Pseudomonas* species detected on dog chin came from the environment since they have been previously isolated from environments such as soil or water sources [39, 40].

The most abundant species in dog dorsal skin samples were *Stenotrophomonas rhizophila*, *Bacillus cereus*, *Sanguibacter keddieii*, *Sporosarcina psychrophila, Achromobacter xylosidans* and *Glutamicibacter arilaitensis*. None of these specific bacterial species had previously been associated with a healthy skin microbiota neither in human nor in dogs. Some of them have an environmental origin such as: *Stenotrophomonas rhizophila*, which is mainly associated to plants [41]; or *Sporosarcina psychrophila,* which is widely distributed in terrestrial and aquatic environments [42]. The *Bacillus cereus* main reservoir is also the soil, although it can be a commensal of root plants and guts of insects and it can also be a pathogen for insects and mammals [43]. Overall, environmental-associated bacteria have already been associated to dog skin microbiota and are expected since dogs constantly interact with the environment [27].

Regarding *Stenotrophomonas* in human microbiota studies, Flores and collaborators found that this genus was enriched in atopic dermatitis patients that were responders with emollient treatment [44]. However, previous studies on this skin disease found *Stenotrophomonas maltophila* associated to the disease rather than *Stenotrophomonas rhizophila* [45]. Also, *Achromobacter xylosoxidans* has been mainly associated to different kind of infections, also skin and soft tissue infections in humans [46]. However, both dogs included in this pool were healthy and with representatives of both genus/species, a fact that reinforces the need to study the healthy skin microbiome before associating some species at the taxonomic level to disease. The other abundant bacteria detected on dog skin have been isolated in very different scenarios: *Sanguibacter keddieii* from cow milk and blood [47, 48]; and *Glutamicibacter arilaitensis* (formerly *Arthrobacter arilaitensis*) is commonly isolated in cheese surfaces [35, 49].

Finally, some of the technical parameters used should be improved for better performance in future studies. One the one hand, in most cases we did not obtain enough DNA mass to begin with the indicated number of molecules for *rrn* operon amplicons. Thus, the flowcell contained an underrepresentation of *rrn* operon amplicons when compared to the full-length *16S rRNA* gene. Moreover, in barcodes that contained *rrn* operon amplicons, a great percentage of reads were lost due to an inaccurate sequence size (∼1,500bp). One possible solution could be running each marker gene in different runs, so multiplexing samples with the same size amplicon to avoid underrepresentation of the larger one. On the other hand, when assessing chimera in mock samples using the specific mock database, we detected that *16S rRNA* gene formed a higher percentage of chimeras than *rrn* operon. Some options to improve that fact would include lowering PCR cycles performed. Better adjusting the laboratory practices would allow an increased DNA yield that meets the first quality control steps.

To conclude, both full-length *16S rRNA* and the *rrn* operon retrieved the microbiota composition from simple and complex microbial communities, even from the low-biomass samples such as dog skin. Taxonomy assignment down to species level was obtained, although it was not always feasible due to: i) sequencing errors; ii) high similarity of the marker chosen within some genera; and iii) incomplete database. For an increased resolution at the species level, *rrn* operon would be the best choice. Further studies should be relying on the ONT 1D^2^ kit, on the new basecallers and in the new flow cells with R10 pores that combined will get higher accuracy. Finally, studies comparing marker-based strategies with metagenomics will determine the most accurate marker for microbiota studies in low-biomass samples.

## Data availability

The datasets analyzed during the current study are available in the SRA NCBI repository under the Bioproject accession number PRJNA495486.

## Authors’ contributions

AS, OF and AC Conceptualization; AC Formal Analysis; CC, JV and AC Investigation; OF, AC Methodology; OF, AS Resources & Funding Acquisition; OF Supervision; CC and AC Validation; CC, JV and AC Visualization; AC Writing – Original Draft Preparation; CC, JV, OF and AC Writing – Review & Editing

## Supplementary material

### Supplementary Table S1. Taxa found on skin microbiota of healthy dogs

List of all the taxa and its relative abundances found on dog skin microbiota samples (chin and dorsal skin). Results for both marker genes tested and for both approaches (Minimap2 + rrn database and WIMP with NCBI database).

## Bibliography

1 Salter SJ, Cox MJ, Turek EM, Calus ST, Cookson WO, Moffatt MF, Turner P, Parkhill J, Loman NJ, Walker AW. Reagent and laboratory contamination can critically impact sequence-based microbiome analyses. BMC Biology (2014). 12(1): 87.

2 Kong HH, Andersson B, Clavel T, Common JE, Jackson SA, Olson ND, Segre JA, Traidl-Hoffmann C. Performing Skin Microbiome Research: A Method to the Madness. J Invest Dermatol (2017). 137(3): 561–568.

3 Ludwig W, Schleifer KH. Bacterial phylogeny based on 16S and 23S rRNA sequence analysis. FEMS Microbiol Rev (1994). 15: 155–173.

4 Yarza P, Ludwig W, Euzéby J, Amann R, Schleifer KH, Glöckner FO, Rosselló-Móra R. Update of the All-Species Living Tree Project based on 16S and 23S rRNA sequence analyses. Syst Appl Microbiol (2010). 33: 291–299.

5 Iwen PC, Hinrichs SH, Ruppy ME. Utilization of the internal transcribed spacer regions as molecular targets to detect and identify human fungal pathogens. Med Mycol (2002). 40: 87–109.

6 Hibbett DS, Ohman A, Glotzer D, Nuhn M, Kirk P, Nilsson RH. Progress in molecular and morphological taxon discovery in Fungi and options for formal classification of environmental sequences. Fungal Biol Rev (2011). 25: 38–47.

7 Clarridge JE. Impact of 16S rRNA Gene Sequence Analysis for Identification of Bacteria on Clinical Microbiology and Infectious Diseases. Clin Microbiol Rev (2004). 17: 840–862.

8 Janda JM, Abbott SL. 16S rRNA gene sequencing for bacterial identification in the diagnostic laboratory: pluses, perils, and pitfalls. J Clin Microbiol (2007). 45: 2761–2764.

9 Walters WA, Caporaso JG, Lauber CL, Berg-Lyons D, Fierer N, Knight R. PrimerProspector: de novo design and taxonomic analysis of barcoded polymerase chain reaction primers. Bioinformatics (2011). 27: 1159–1161.

10 Kuczynski J, Lauber CL, Walters WA, Parfrey LW, Clemente JC, Gevers D, Knight R. Experimental and analytical tools for studying the human microbiome. Nat Rev Genet (2012). 13: 47–58.

11 Grice EA. The skin microbiome: potential for novel diagnostic and therapeutic approaches to cutaneous disease. Semin Cutan Med Surg(2014). 33: 98–103.

12 Chng KR, Tay ASL, Li C, Ng AHQ, Wang J, Suri BK, Matta SA, McGovern N, Janela B, Wong XF, et al. Whole metagenome profiling reveals skin microbiome-dependent susceptibility to atopic dermatitis flare. Nat Microbiol (2016) 1:16106

13 Pierezan F, Olivry T, Paps JS, Lawhon SD, Wu J, Steiner JM, Suchodolski JS, Hoffmann AR. The skin microbiome in allergen-induced canine atopic dermatitis. Vet dermatol(2016). 27:332-e82.

14 Bradley CW, Morris DO, Rankin SC, Cain CL, Misic AM, Houser T, Mauldin EA, Grice EA. Longitudinal evaluation of the skin microbiome and association with microenvironment and treatment in canine atopic dermatitis. J Invest Dermatol (2016). 136: 1182–1190.

15 Li C, Chng KR, Boey EJ, Ng AH, Wilm A, Nagarajan N. INC-Seq: accurate single molecule reads using nanopore sequencing. Gigascience (2016). 5:34.

16 Benítez-Páez A, Portune KJ, Sanz Y. Species-level resolution of 16S rRNA gene amplicons sequenced through the MinION^TM^ portable nanopore sequencer. Gigascience (2016). 5:4.

17 Brown BL, Watson M, Minot SS, Rivera MC, Franklin RB. MinION^TM^ nanopore sequencing of environmental metagenomes: a synthetic approach. Gigascience (2017). 6:1–10.

18 Shin J, Lee S, Go MJ, Lee SY, Kim SC, Lee CH, Cho BK. Analysis of the mouse gut microbiome using full-length 16S rRNA amplicon sequencing. Sci Rep (2016). 6:29681.

19 Ma X, Stachler E, Bibby K. Evaluation of Oxford Nanopore MinION Sequencing for 16S rRNA Microbiome Characterization. bioRxiv (2017).

20 Shin H, Lee E, Shin J, Ko SR, Oh HS, Ahn CY, Oh HM, Cho BK, Cho S. Elucidation of the bacterial communities associated with the harmful microalgae Alexandrium tamarense and Cochlodinium polykrikoides using nanopore sequencing. Sci Rep (2018). 8:5323.

21 Cusco A, Vines J, D’Andreano S, Riva F, Casellas J, Sanchez A, Francino O. Using MinION to characterize dog skin microbiota through full-length 16S rRNA gene sequencing approach. bioRxiv (2017).

22 Mitsuhashi S, Kryukov K, Nakagawa S, Takeuchi JS, Shiraishi Y, Asano K, Imanishi T. A portable system for rapid bacterial composition analysis using a nanopore-based sequencer and laptop computer. Sci Rep (2017). 7:5657.

23 Moon J, Kim N, Lee HS, Shin HR, Lee ST, Jung KH, Park KI, Lee SK, Chu K. Campylobacter fetus meningitis confirmed by a 16S rRNA gene analysis using the MinION nanopore sequencer, South Korea, 2016. Emerg Microbes Infect (2017). 6:e94.

24 Moon J, Jang Y, Kim N, Park WB, Park KI, Lee ST, Jung KH, Kim M, Lee SK, Chu K. Diagnosis of Haemophilus influenzae Pneumonia by Nanopore 16S Amplicon Sequencing of Sputum. Emerg Infect Dis (2018). 24: 1944–1946.

25 Benítez-Páez A, Sanz Y. Multi-locus and long amplicon sequencing approach to study microbial diversity at species level using the MinION^TM^ portable nanopore sequencer. Gigascience (2017). 6: 1–12.

26 Kerkhof LJ, Dillon KP, Häggblom MM, McGuinness LR. Profiling bacterial communities by MinION sequencing of ribosomal operons. Microbiome (2017). 5:116.

27 Cuscó A, Belanger JM, Gershony L, Islas-Trejo A, Levy K, Medrano JF, Sánchez A, Oberbauer AM, Francino O. Individual Signatures and Environmental Factors Shape Skin Microbiota on Healthy Dogs. Microbiome (2017). 5:139.

28 Zeng YH, Koblížek M, Li YX, Liu YP, Feng FY, Ji JD, Jian JC, Wu ZH. Long PCR-RFLP of 16S-ITS-23S rRNA genes: a high-resolution molecular tool for bacterial genotyping. J Appl Microbiol (2013). 114: 433–447.

29 Klindworth A, Pruesse E, Schweer T, Peplies J, Quast C, Horn M, Glöckner FO. Evaluation of general 16S ribosomal RNA gene PCR primers for classical and next-generation sequencing-based diversity studies. Nucleic Acids Res (2013), 41:e1.

30 Wick R. Porechop. Available at: https://github.com/rrwick/Porechop

31 Li H. Minimap2: pairwise alignment for nucleotide sequences. Bioinformatics (2018), 34: 3094–3100.

32 Marijon P. yacrd: Yet Another Chimeric Read Detector for long reads. Available at: https://github.com/natir/yacrd

33 Juul S, Izquierdo F, Hurst A, Dai X, Wright A, Kulesha E, Pettett R, Turner DJ. What’s in my pot? Real-time species identification on the MinION. bioRxiv (2015).

34 Kim D, Song L, Breitwieser FP, Salzberg SL. Centrifuge: rapid and sensitive classification of metagenomic sequences. Genome Res (2016). 26: 1721–1729.

35 Busse HJ. Review of the taxonomy of the genus Arthrobacter, emendation of the genus Arthrobacter sensu lato, proposal to reclassify selected species of the genus Arthrobacter in the novel genera Glutamicibacter gen. nov., Paeniglutamicibacter gen. nov., Pseudoglutamicibacter gen. nov., Paenarthrobacter gen. nov. and Pseudarthrobacter gen. nov., and emended description of Arthrobacter roseus. Int J Syst Evol Microbiol (2016). 66: 9–37.

36 Ghebremedhin B, Layer F, König W, König B. Genetic classification and distinguishing of Staphylococcus species based on different partial gap, 16S rRNA, hsp60, rpoB, sodA, and tuf gene sequences. J Clin Microbiol (2008). 46: 1019–1025.

37 Meason-Smith C, Older CE, Ocana R, Dominguez B, Lawhon SD, Wu J, Patterson AP, Rodrigues Hoffmann A. Novel association of Psychrobacter and Pseudomonas with malodour in bloodhound dogs, and the effects of a topical product composed of essential oils and plant-derived essential fatty acids in a randomized, blinded, placebo-controlled study. Vet Dermatol (2018).

38 Riggio MP, Lennon A, Taylor DJ, Bennett D. Molecular identification of bacteria associated with canine periodontal disease. Vet Microbiol (2011). 150: 394–400.

39 Peix A, Ramírez-Bahena MH, Velázquez E. Historical evolution and current status of the taxonomy of genus Pseudomonas. Infect Genet Evol (2009). 9: 1132–1147.

40 Mehri I, Turki Y, Chair M, Chérif H, Hassen A, Meyer JM, Gtari M. Genetic and functional heterogeneities among fluorescent Pseudomonas isolated from environmental samples. J Gen Appl Microbiol (2011). 57: 101–114.

41 Wolf A, Fritze A, Hagemann M, Berg G. Stenotrophomonas rhizophila sp. nov., a novel plant-associated bacterium with antifungal properties. Int J Syst Evol Microbiol (2002). 52: 1937–1944.

42 Yan W, Xiao X, Zhang Y. Complete genome sequence of the Sporosarcina psychrophila DSM 6497, a psychrophilic Bacillus strain that mediates the calcium carbonate precipitation. J Biotechnol (2016). 226: 14–15.

43 Ceuppens S, Boon N, Uyttendaele M. Diversity of Bacillus cereus group strains is reflected in their broad range of pathogenicity and diverse ecological lifestyles. FEMS Microbiol Ecol (2013). 84: 433–450.

44 Seite S, Flores GE, Henley JB, Martin R, Zelenkova H, Aguilar L, Fierer N. Microbiome of affected and unaffected skin of patients with atopic dermatitis before and after emollient treatment. J Drugs Dermatol (2014). 13: 1365–1372.

45 Dekio I, Sakamoto M, Hayashi H, Amagai M, Suematsu M, Benno Y. Characterization of skin microbiota in patients with atopic dermatitis and in normal subjects using 16S rRNA gene-based comprehensive analysis. J Med Microbiol (2007). 56: 1675–1683.

46 Tena D, Martínez NM, Losa C, Solís S. Skin and soft tissue infection caused by Achromobacter xylosoxidans: report of 14 cases. Scand J Infect Dis (2014). 46: 130–135.

47 Fernández-Garayzábal JF, Dominguez L, Pascual C, Jones D, Collins MD. Phenotypic and phylogenetic characterization of some unknown coryneform bacteria isolated from bovine blood and milk: description of Sanguibacter gen.nov. Lett Appl Microbiol (1995). 20: 69–75.

48 Ivanova N, Sikorski J, Sims D, Brettin T, Detter JC, Han C, Lapidus A, Copeland A, Glavina Del Rio T, Nolan M, et al. Complete genome sequence of Sanguibacter keddieii type strain (ST-74). Stand Genomic Sci (2009). 1: 110–118.

49 Irlinger F, Bimet F, Delettre J, Lefèvre M, Grimont PA. Arthrobacter bergerei sp. nov. and Arthrobacter arilaitensis sp. nov., novel coryneform species isolated from the surfaces of cheeses. Int J Syst Evol Microbiol (2005). 55: 457–462.

